# Dynamic expression of Id3 defines the stepwise differentiation of tissue-resident regulatory T cells

**DOI:** 10.1101/359687

**Authors:** Jenna M. Sullivan, Barbara Höllbacher, Daniel J. Campbell

**Author notes:** Corresponding Author: Daniel J. Campbell, Benaroya Research Institute, 1201 Ninth Avenue, Seattle, WA 98101-2795.

## Abstract

Foxp3^+^ regulatory T (T_R_) cells are phenotypically and functionally diverse, and broadly distributed in lymphoid and non-lymphoid tissues. However, the pathways guiding the differentiation of tissue-resident T_R_ populations have not been well defined. By regulating E-protein function, Id3 controls the differentiation of CD8^+^ effector T cells and is essential for T_R_ maintenance and function. We show that dynamic expression of Id3 helps define three distinct T_R_ populations, Id3^+^CD62L^hi^CD44^lo^ central (c)T_R,_ Id3^+^CD62L^lo^CD44^hi^ effector (e)T_R_ and Id3^-^ eT_R_. Adoptive transfer experiments and transcriptome analyses support a stepwise model of differentiation from Id3^+^ cT_R_ to Id3^+^ eT_R_ to Id3^-^ eT_R_. Furthermore, Id3^-^ eT_R_ have high expression of functional inhibitory markers and a transcriptional signature of tissue-resident T_R_. Accordingl Id3^-^ eT_R_ are highly enriched in non-lymphoid organs, but virtually absent from blood and lymph Thus, we propose that tissue-resident T_R_ develop in a multi-step process associated with Id3 downregulation.

## Introduction

Several recent studies have highlighted the phenotypic and functional heterogeneity of regulatory T (T_R_) cells during both steady state and inflammation (1-4). We and others have shown that at steady state in lymphoid organs T_R_ can be broadly divided by expression of CD4 and CD62L into distinct subsets which differ in their localization, dependence on IL-2, and extent of PI3K signaling (2, 5, 6). Moreover, CD44^hi^CD62L^lo^ effector (e)T_R_ display diverse expression of transcription factors and chemokine receptors that promote their migration to inflamed tissues and their response to different types of inflammatory signals (1, 7). Accordingly, T_R_ found in nonlymphoid tissues have a distinct molecular profile that includes high expression of Gata3 and ST2 (the IL-33R), and are functionally equipped to suppress inflammation at barrier sites (8, 9). Although these data highlight the anatomical, functional and molecular diversity of T_R_, the pathways by which these T_R_ populations differentiate have not been completely defined.

The inhibitors of DNA binding (Id) proteins have been extensively studied in lymphocyte development (10, 11). Studies of CD8^+^ effector T cells revealed that Id2 and Id3 are powerful transcriptional regulators of differentiation that are dynamically regulated during T cell activation and effector/memory T cell differentiation (12, 13). Through their regulation of E protein function Id2 and Id3 help to control expression of genes essential for CD8^+^ effector cell differentiation and survival such as *Tcf7*, *Tbx21*, *Bcl2* and *Klrg1* (14, 15). Although less well studied, Id proteins have been shown to have essential roles in CD4^+^ T cell function. For instance, Id2 and Id3 are essential for T_R_ maintenance and function, with T_R_ lacking both Id2 and Id3 having impaired proliferation and survival (16). In TR, Id3 helps to stabilize Foxp3 through restriction of the E protein E47 and its downstream targets Spi-B and SOCS3 (17). However, Id3 expression is not uniform in TR, and distinct populations of Id3^+^ and Id3^-^ have been identified (16, 18). In this study, we show that Id3 is dynamically regulated in TR, and that progressive loss of Id3 correlates with the stepwise differentiation of a highly-functional T_R_ population localized primar in non-lymphoid tissues.

## Materials and Methods

### Mice

C57BL/6, RAG1-deficient and Foxp3-mRFP mice were purchased from The Jackson Laboratory. Id3-GFP mice were a gift from Ananda Goldrath (UCSD, La Jolla, California) and have been previously described (12, 14). Mice were bred and housed under the approval of the Institutional Animal Care and Use Committee of the Benaroya Research Institute.

### Cell Isolation

Unless noted below, single cell suspensions isolated from tissues using manual disruption. PEC isolated by injecting sterile PBS into peritoneal cavity of euthanized mice, gentle disruption to dislodge cells and collection of injected PBS. IEL and LPL were isolated from pooled large and small intestine as previous described (19). Lymphocytes further purified by resuspension in 44% Percoll^TM^ (GE Healthcare) layered over 67% Percoll^TM^ and spun at 2,800rpm for 20 mins. Lung and fat were finely minced, digested with 0.26U/mL Liberase TM (Roche) and 10U/mL DNAse (Sigma) for 1 hr at 37°C and filtered. For skin tissue, ears were processed as above with 0.14U/mL Liberase TM and 10U/mL DNAse. For lymph collection, mice were fed 20mL/kg ‘Half and Half’ by oral gavage and sacrificed 2-3 hrs later. Lymph collected from the cisterna chyli was directly stained for flow cytometry.

### Flow cytometry

Single cells suspensions were stained with fixable Viability Dye eFluor 780 (eBiosciences) in PBS for 10 min at RT. Cells were stained with directly conjugated Abs in PBS with 0.5% BCS for 20 min at 4°C. Abs purchased from BioLegend: CD4 (RM4-5), TCRβ (H57-597), CD44 (IM7), CD62L (MEL-14), CD25 (PC61), ICOS (C398.4A), TIGIT (1G9), CTLA4 (UC10-4B9), GITR (DTA.1), CD69 (53-7.3). Abs purchased from eBiosciences: KLRG1 (2F1) and CD103 (2E7). Intracellular stains were performed using a FixPerm Kit (eBiosciences). Data acquired on an LSR II (BD Biosciences) and analyzed using FlowJo software (TreeStar).

### *In vitro* assays

CD4^+^ T cells isolated from spleen and LN using CD4 microbeads (Miltenyi). 1×10^6^ T cells cultured with platebound α-CD3 (2C11) and α-CD28 (37.51) from BioXcell at 1μg/mL each for 48 or 66 hrs. Inhibitors purchased and used as follows: ZSTK474 (1μM, Sigma), Rapamycin (10nM, Selleckchem), NFAT inhibitor (10μM, Tocoris), Mek inhibitor PD0325901 (100nM, Peprotech) and Erk inhibitor FR180204 (10μM, Tocoris). *In vitro* T_R_ suppression assays were performed as previous described (20). Chemotaxis assay performed as previously described (21).

### *In vivo* T_R_ transfer

Sorted T_R_ isolated from spleen and LN as described for RNA-seq. 100,000 sorted cells were injected retro-orbitally into RAG1-deficient hosts. Spleen, LN and blood of recipient mice collected two wks later and analyzed by flow cytometry.

### Statistical Analysis

The *p* values were calculated by Prism software (GraphPad) using either an unpaired Student’ *t* test or one way ANOVA as indicated. Values less than 0.05 were considered significant.

### RNA-seq

CD4^+^ T cells were isolated using CD4 microbreads (Miltenyi) from either peripheral LNs or spleens of 3 littermate Id3-GFP × Foxp3-mRFP mice. Cells were sorted based on viability, CD4, CD44, CD62L, Id3-GFP and Foxp3-mRFP expression on a FACs Aria II (BD Biosciences). 500 cells were sorted directly into SMART-Seq v4 Ultra Low Input RNA Kit (Takara) lysis buffer and protocol followed to produce cDNA. Library construction was performed using a modified protocol of the NexteraXT DNA sample preparation kit (Illumina). Dual-index, single-read sequencing of pooled libraries was run on a HiSeq2500 sequencer (Illumina) with 58-base reads and an average depth of 8.6Mio reads per library. Base-calling and demultiplexing were performed automatically on BaseSpace (Illumina) to generate FASTQ files.

## Results/Discussion

### Id3 is dynamically expressed in T_R_ and regulated by TCR signaling

To examine Id3 expression in T_R_ we generated Id3-GFP × Foxp3-mRFP double reporter mice. In agreement with previously published reports, most CD4^+^ T cells in spleen and lymph nodes (LNs) were Id3^+^ (16, 18). However, we noted a subset of Id3^-^ cells in both Foxp3^+^ T_R_ and Foxp3^-^ conventional CD4^+^ T cell populations (Fig 1A). Within T_R_, the Id3^-^ population fell exclusively within the CD44^hi^ CD62L^lo^ eT_R_ compartment (5), whereas Id3^+^ T_R_ were found in both the CD44^lo^ CD62L^hi^ central (c)T_R_ and eT_R_ compartments (Fig 1B). Thus, in secondary lymphoid organs (SLOs) T_R_ can be divided into three distinct subsets based on Id3, CD62L and CD44 expression, Id3^+^ cT_R_, Id3^+^ eT_R_ or Id3^-^ eT_R_. Within the Id3^+^ T_R_ populations Id3^+^ cT_R_ had higher Id3 expression than Id3^+^ eT_R_ measured by GFP-MFI (Fig 1B), leading us to hypothesize that T may downregulate Id3 as they transition during their activation and differentiation from cT_R_ into eT_R_. Indeed, Id3 expression can be downregulated by TCR signaling (16), and we observed a loss of Id3 expression in the T_R_ compartment correlating with the strength of TCR stimulation when cells were activated with different concentrations of platebound αCD3/28 (Sup Fig 1). Furthermore, inhibiting either the MAP-kinase/Erk or PI3-kinase/mTOR signaling pathways blocked Id3 downregulation in T_R_ (Fig 1C), consistent with reports that eT_R_ development requires TCR stimulation, mTOR signaling and the PI3-kinase-dependent inactivation of the transcription factor Foxo1 (5, 6, 22).

**Figure 1:**
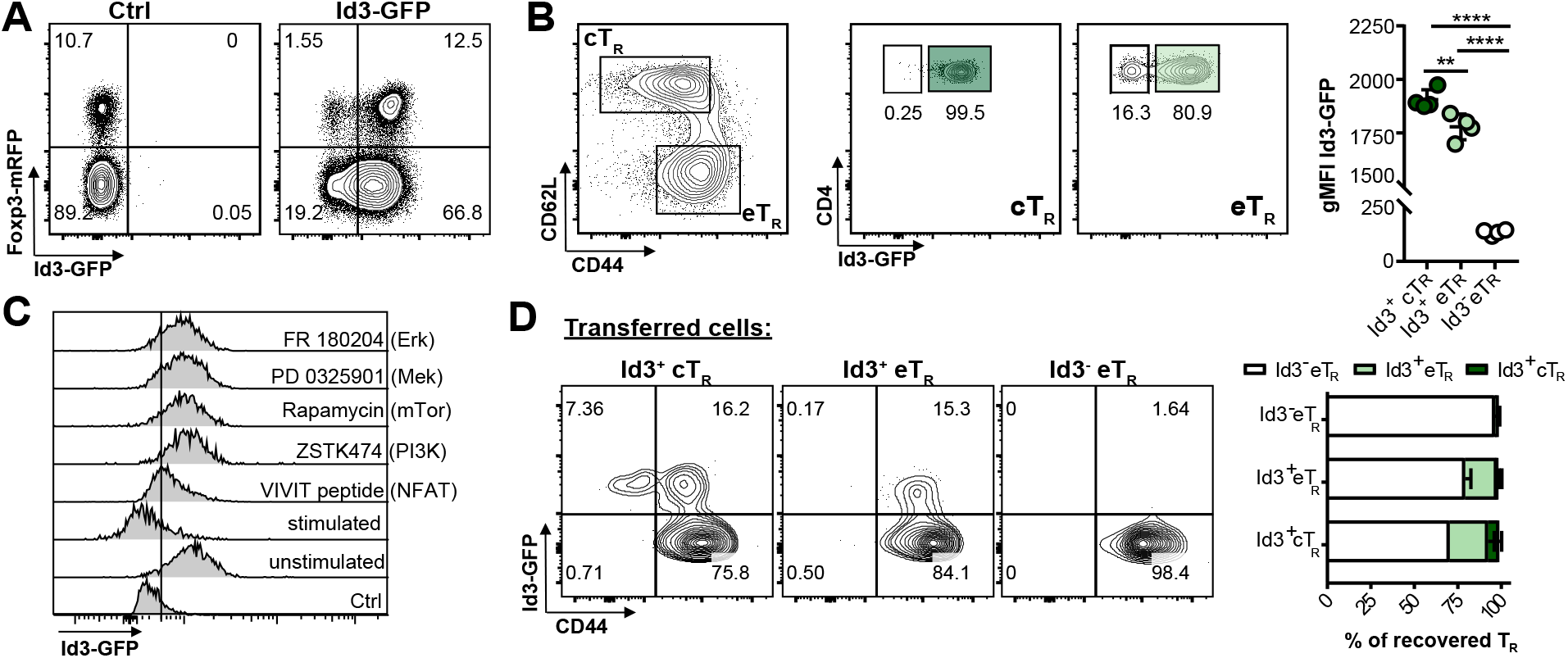
Id3 is dynamically expressed in T_R_ and regulated by TCR signaling. A) Representative flow cytometry plots of Id3-GFP and Foxp3-RFP expression by gated spleni TCRβ^+^ CD4^+^ T cells. B) Representative flow cytometry analysis of Id3-GFP expression by splenic CD44^lo^CD62L^hi^ cT_R_ and CD44^hi^CD62L^lo^ eT_R_ gated as indicated. (Bottom left) Graphical analysis of gMFI of Id3-GFP in each of the three gated T_R_ populations. C) Representative flow cytometry plots of Id3-GFP expression by gated splenic TCRβ^+^ CD4^+^ Foxp3^+^ T_R_ 66 hrs after stimulation of purified CD4^+^ T cells in the presence or absence of the idicated inhibitors, representative of 2 independent expts. D) Representative flow cytometry plots and graphical analysis of CD44 and Id3-GFP expression by gated TCRβ^+^CD4^+^Foxp3^+^ T_R_ recovered from the LNs of RAG1-deficient mice 2 weeks after transfer of the indicated T_R_ population, summary of 3 independent expts, 3-5 mice per group total. Significance determined by one way ANOVA with Tukey’s post-test for pairwise comparisons, ^*^p <0.05, ^**^p < 0.01, ^****^p < 0.001

To more precisely define the developmental relationship between these T_R_ subsets we utilized an adaptive transfer model in which T_R_ stimulation and expansion depends on TCR:MHCII interactions (23). For this we sorted Id3^+^ cTR, Id3^+^ eT_R_ or Id3^-^ eT_R_ from spleen and LNs of reporter mice and transferred individual TR populations into RAG1-deficient animals, and evaluated the phenotype and expansion of transferred T_R_ after 2 weeks. Importantly, we did not observe any difference in the extent of Foxp3 expression between the T_R_ populations upon their recovery, which varied between ~30-80% in different experiments (not shown). Transferred Id3^+^ cT_R_ gave rise to all three subsets, with some cells retaining Id3 and CD62L expression but the majority converting into Id3^-^ eT_R_ (Fig 1D). The bulk of Id3^+^ eT_R_ downregulated Id3, with no cells regaining CD62L expression, whereas Id3^-^ eT_R_ did not give rise to either of the other populations, indicating that these cells are likely a terminal differentiated population. Thus, Id3^-^ eT_R_ appear to develop from Id3^+^ cT_R_ in a stepwise manner during activation, with Id3^+^ eT_R_ acting as an intermediate population.

### Id3^-^ eT_R_ express inhibitory markers and are highly suppressive

To explore the phenotypic and functional differences between these three subsets of T_R_, we assessed expression of the T_R_-associated surface markers on each population of splenic T_R_. In agreement with previously published data, cT_R_ and eT_R_ showed distinct phenotypes, with eT_R_ having lower expression of the high affinity IL-2 receptor component CD25, but higher levels of the activation and functional surface markers ICOS, KLRG1, TIGIT, GITR and CTLA4 (Fig 2A) (5, 6). Moreover, within the eT_R_ compartment the Id3^-^ T_R_ had higher expression of activation and inhibitory molecules than their Id3^+^ T_R_ counterparts, but had the lowest expression of CD25. As our lab previously described (5), elevated expression of ICOS and diminished expression of CD25 suggests that Id3^-^ eT_R_ are less dependent on IL-2 for their homeostatic maintenance, an instead likely rely on continued ICOS signaling. Additionally, increased expression of these T_R_ functional molecules correlated with enhanced *in vitro* suppressive activity of Id3^-^ eT_R_ compare with either Id3^+^ cT_R_ or Id3^+^ eT_R_ (Fig 2B).

**Figure 2:**
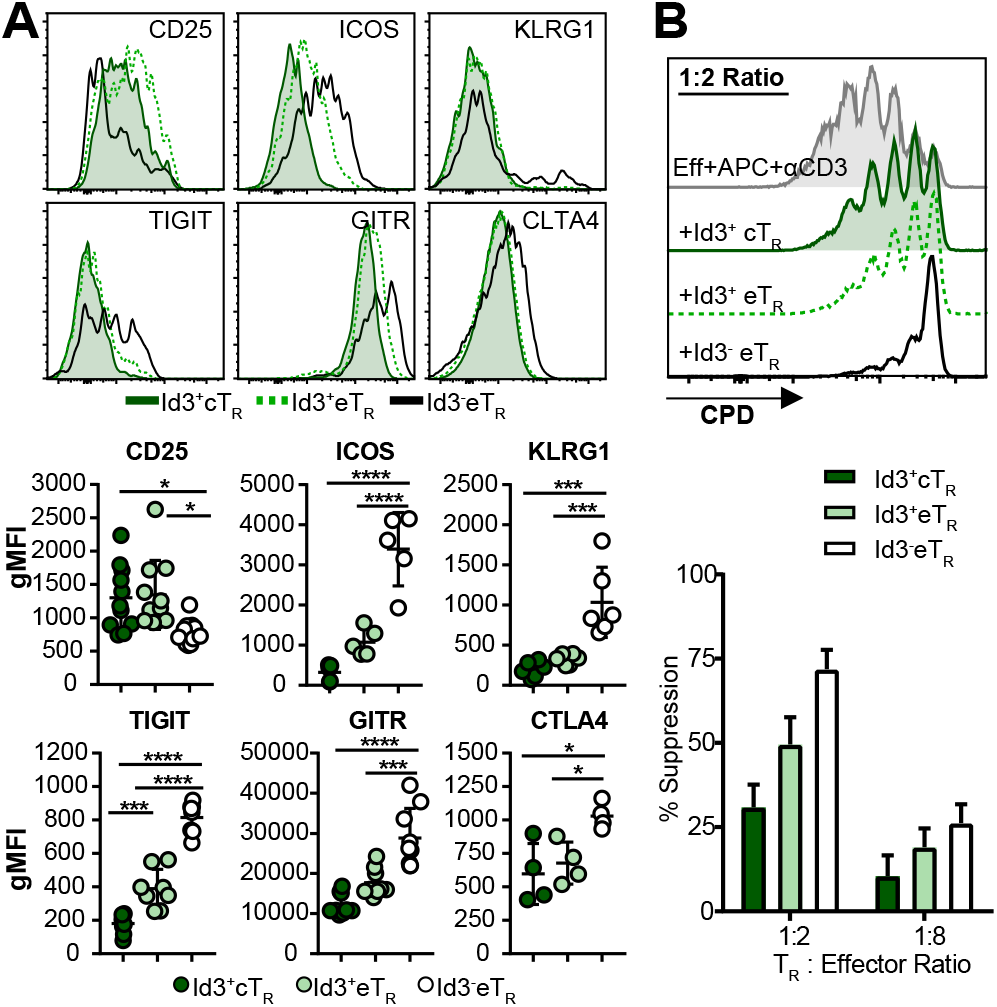
Id3^-^ eT_R_ express inhibitory markers and are highly suppressive. A) (Top) Representative flow cytometry histograms. (Bottom) Graphical analysis of expression of the indicated markers by gated splenic T_R_ populations. B) (Top) Representative flow cytometry analysis of CPD dilution by CD4^+^Foxp3^-^ effector T cells stimulated with or without the indicated T_R_ populations. (Bottom) Graphical analysis of suppression by each of the indicated populations, n=3. Significance determined by one way ANOVA with Tukey’s post-test for pairwise comparisons, ^*^p <0.05, ^**^p < 0.01, ^****^p < 0.001

### Transcriptional profiling highlights the stepwise differentiation of Id3^-^ eT_R_

To identify and compare their unique transcriptional profiles, we performed RNA-seq on sorted Id3^+^ cT_R_, Id3^+^ eT_R_ and Id3^-^ eT_R_ from spleen or LNs of Id3-GFP × Foxp3-mRFP reporter mice. Although there was little difference between LN and spleen samples, principle component analysis (PCA) showed that each of the three T_R_ populations were transcriptionally distinct (Fig 3A), and accordingly we identified 1,672 significantly differentially expressed (DE) genes between the three T_R_ populations (Fig 3B, Supplemental Table 1). The largest differences were found between Id3^-^ eT_R_ and Id3^+^ cT_R_ with 1,471 DE genes, whereas only 474 genes differed between Id3^+^ and Id3^-^ eT_R_ (Fig 3B). Interestingly among the DE genes we observed a reciproc increase in Id2 expression as T_R_ lose Id3 (Fig 3C). This differential Id expression in T_R_ is simila to what occurs in CD8^+^ T cells, which upregulate Id2 and downregulate Id3 while becoming activated and gaining effector function (12, 13). This suggests that as T_R_ downregulate Id3, the closely related Id2 may take over some of its functions while also driving a unique E-protein-dependent signature promoting eT_R_ development. Consistent with the stepwise differentiation model we propose for these populations, examination of the 300 most DE genes across the three T_R_ subsets (based on highest F-value) showed that both up and downregulated genes were generally expressed in a gradient fashion, with expression in Id3^+^ eT_R_ falling between that of Id3^+^ cT_R_ and Id3^-^ eT_R_ (Fig 3D-E). Moreover, in accordance with our prior phenotypic analysis and their enhanced suppressive activity, Id3^-^ eT_R_ showed elevated expression of known T_R_ function genes, including *Il10, Ctla4, Pdcd1, Tnfrsf4, Lag3 and Ebi3* (Fig 3F). To further validate our RNA-seq results, we confirmed differential expression of several T_R_-associated genes by flow cytometry (Supplemental Fig 2). Gene Ontology (GO) term enrichment analysis of genes DE between Id3^+^ eT_R_ and Id3^-^ eT_R_ identified specific molecular pathways altered between these closely related populations (Fig 3G). The top six enriched categories all related to cytokine and chemokine receptor signaling, with Id3^-^ eT_R_ expressing high levels of receptors indicating that they can tune their activity in response to key inflammatory cytokines such as IL-1, IL-18, IL-23, and IL-25 (Fig 3H). Moreover, among the DE chemokine receptors, Id3^-^ eT_R_ had the highest expression and subsequent responsiveness to chemokines that promote lymphocyte migration to inflamed tissues, such as *Ccr4* and *Cxcr3* (Fig 3H-I) (1). Thus, Id3^-^ eT_R_ have a unique molecular profile indicative of their development from Id3^+^ T_R_ precursors, their elevated suppressive function and altered migratory capacity.

**Figure 3:**
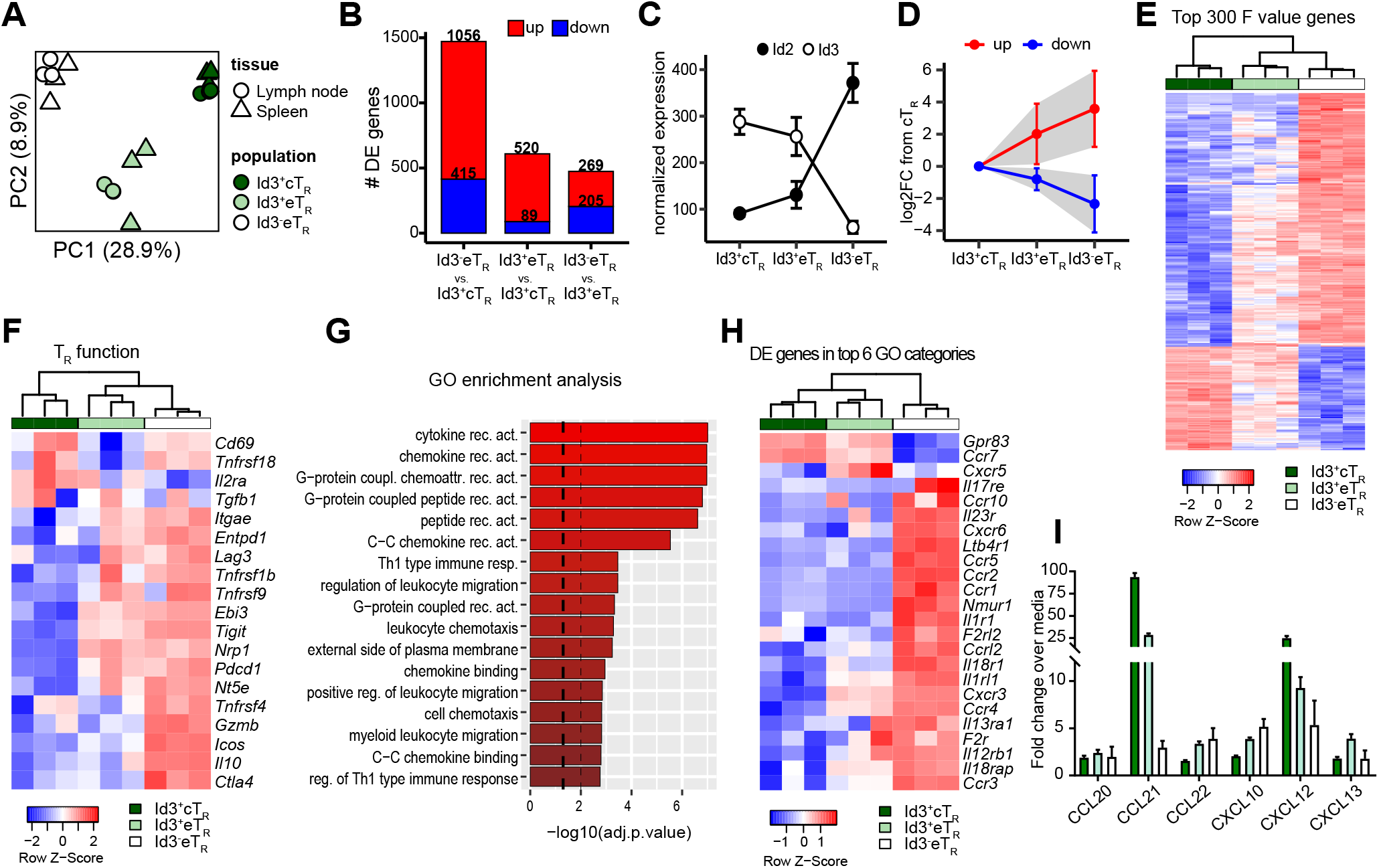
Transcriptional profiling highlights the stepwise differentiation of Id3^-^ eT_R_. A) PCA of RNA-seq data from LN and splenic Id3^+^ cT_R_, Id3^+^ eT_R_ and Id3^-^ eT_R_ populations sorted from three individual mice. B) Bar graphs showing the number of differentially expressed genes (adj.p.value < 0.05 and log2FC >1) for each of the indicated pairwise comparisons. C) Graphical analysis of normalized transcript reads for Id2 or Id3 from RNA-seq data. D) The 300 most differentially expressed genes (determined by F value) were split into the upregulated (red) an downregulated (blue) fractions based expression in Id3^-^ eT_R_ vs Id3^+^ cT_R_. Graph shows the mean log2FC compared to Id3^+^ cT_R_ for both eT_R_ populations. Error bars and shaded area represent 1× SD. E) Heatmap and hierarchical clustering of splenic RNA-seq samples based on the 300 most variably expressed genes. F) Heatmap and hierarchical clustering of splenic RNA-seq samples based on T_R_ signature genes identified in reference (8). G) GO term enrichment analysis for DE genes of Id3^-^ eT_R_ vs. Id3^+^ eT_R_. Dashed lines represent adjusted p values of 0.05 and 0.01. H) Heatmap and hierarchical clustering of splenic RNA-seq samples based on DE genes found in the top 6 GO functional categories enriched in the comparison of Id3^+^ and Id3^-^ eT_R_. I) Graphical analysis of chemotaxis assay.

### Id3^-^ T_R_ are enriched and resident in non-lymphoid tissues

In contrast to their elevated expression of inflammatory chemokine receptors, Id3^-^ eT_R_ had the lowest expression of *Ccr7* and *S1pr1* which function together to promote T_R_ recirculation through SLOs (24) (Fig 3H). Additionally, expression of CD103 and CD69, which together act to retain tissue-resident memory T(_RM_) cells in non-lymphoid sites, was strongly enriched in Id3^-^ eT_R_, suggesting these cells may be tissue-resident (Fig 4A) (25, 26). Indeed, utilizing published gene signatures of CD8^+^ T_RM_ or circulating memory T cells (27), we found that the CD8^+^ T_RM_ signature gene set was enriched in Id3^-^ eT_R_ compared to either Id3^+^ cT_R_ or Id3^+^ eT_R_, whereas the circulating memory gene set was enriched in the Id3^+^ T_R_ populations (Fig 4B). Several groups have recently identified the ST2 (IL-33R)-Gata3 axis as a key determinant of T_R_ residency and function in non-lymphoid tissues (8, 9, 28). Accordingly, Id3^-^ eT_R_ had the highest expression of all positively associated tissue T_R_ genes such as *Il1rl1* (ST2), *Gata3*, *Areg*, *Irf4 and Rora*, whereas Id3^+^ cT_R_ had the lowest expression of these genes but high expression of negative regulators of tissue residence such as *Tcf7, Klf2* and *Lef1,* and Id3^+^ eT displayed intermediate expression for all of these genes (Fig 4C). Consistent with their T_RM_-like transcriptional signature, Id3^-^ eT_R_ were a minority of T_R_ in LNs and spleen, but their frequency dramatically increased in nonlymphoid tissues, where they highly expressed the T_RM_ surface markers CD103 and CD69 (Fig 4D-F). Indeed, tissues such as the fat and skin contained almost exclusively Id3^-^ eT_R_. Of particular note Id3^-^ eT_R_ were rarest in the lymph and blood, indicating that Id3^-^ eT_R_ do not actively recirculate, but instead are retained as tissue-resident cells in non-lymphoid organs.

**Figure 4:**
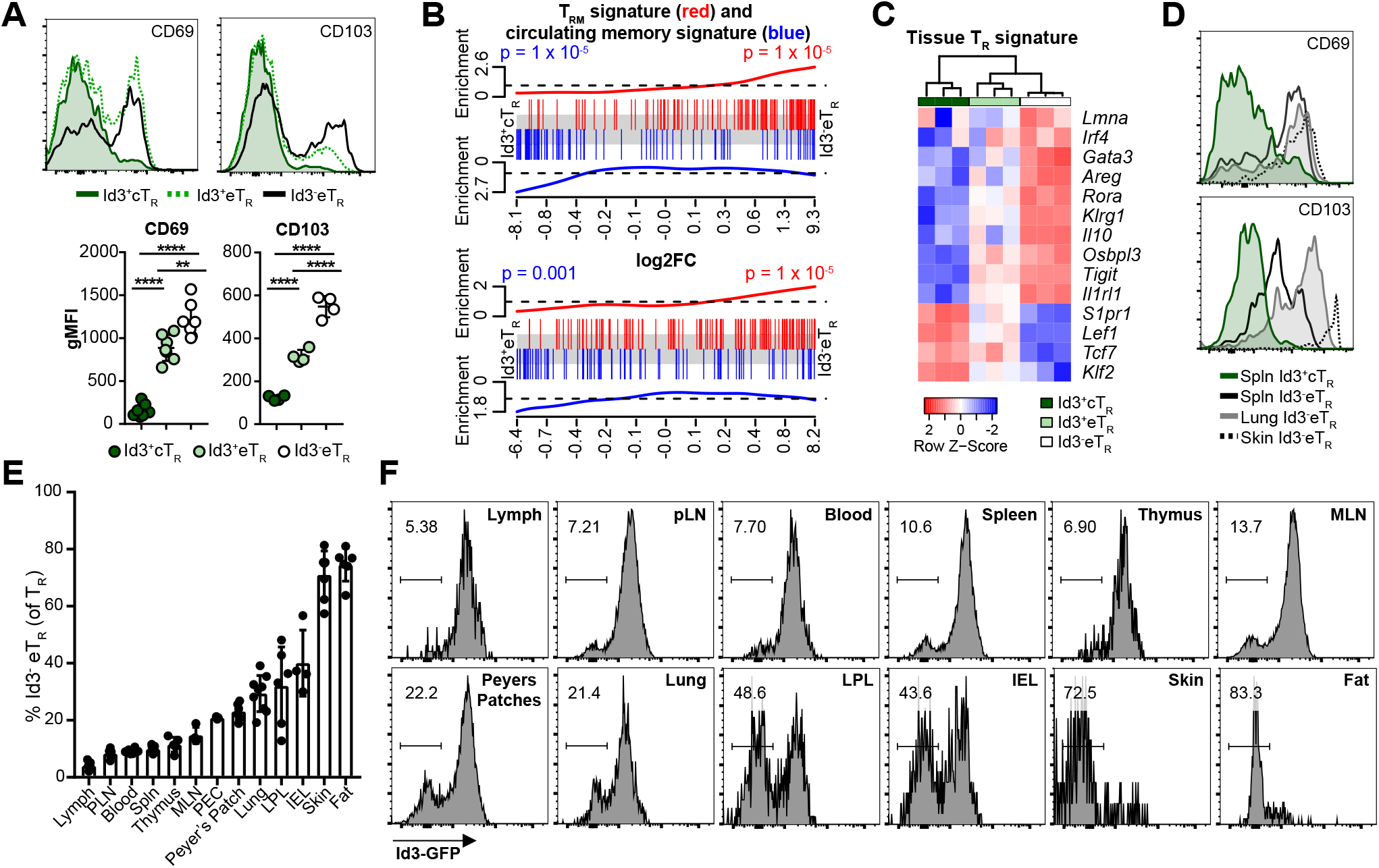
Id3^-^ eT_R_ are enriched and resident in non-lymphoid tissues. A) Representative flow cytometry histograms and graphical analysis of expression of the CD69 and CD103 by gated splenic T_R_ populations. B) Enrichment of CD8^+^ T_RM_ (red) or circulating T cell (blue) gene sets along ranked lists of the indicated pairwise comparisons. C) Heatmap and hierarchical clustering of splenic RNA-seq samples based on ST2^+^ tissue T_R_ associated genes D) Representative flow cytometry plots of CD103 and CD69 expression by gated TCRβ^+^ CD4^+^ Foxp3^+^ T_R_ in the indicated tissues. E) Graphical analysis of Id3^-^ eT_R_ frequency among total T_R_ in various tissues. F) Representative histograms of GFP expression from various tissues of Id3-GFP × Foxp3-mRFP mice, gated on TCRβ^+^ CD4^+^ Foxp3^+^ T_R_.

Despite significant interest in tissue-resident T_R_, the mechanisms regulating their differentiation and distribution are still poorly defined. Our data show that T_R_ can be subdivided based on Id3 expression and known markers of cT_R_ and eT_R_ into distinct populations, and together our phenotypic, functional, transcriptional and transfer analyses strongly support a stepwise differentiation model in which T_R_ downregulate Id3 as they progressively gain effector function, tissue homing capacity and residency in non-lymphoid organs. Similarly, Li *et al.* recently proposed a stepwise model for the differentiation of T_R_ in adipose tissue in which activation in the spleen allowed T_R_ to migrate into the adipose tissue, where IL-33 signaling drove their terminal differentiation and functional specialization (28). High expression of co-stimulatory receptors such as ICOS and receptors for inflammatory cytokines would also allow Id3^-^ eT_R_ to respond to inflammatory signals that enhance Foxp3-mediated transcriptional repression and promote eT_R_ differentiation and function (29), in part through activation of mTORC1 signaling (22). Although the direct role of Id3 downregulation in the functional differentiation of T_R_ is still not established, our data highlight the complexity of tissue T_R_ development, and identify novel molecular pathways that are modulated during their stepwise differentiation.

## Acknowledgements

We thank A. Goldrath for Id3-GFP mice. K. Arumuganathan and T. Nguyen for help with flow cytometry and cell sorting. V. Gersuk and the BRI Genomics Core for running RNA-seq samples. And members of the Campbell lab for helpful discussions.

---

<u>Supplementary Figure 1: Loss of Id3 expression correlates with strength of TCR signaling</u>

Representative flow cytometry plots of Id3-GFP expression by gated splenic TCRβ^+^ CD4^+^ Foxp3^+^ TR 48 hours after stimulation of purified CD4^+^ T cells with 1μg/mL α-CD28 and indicated titration of α-CD3, summary of 2 independent expts.

<u>Supplementary Figure 2: Flow cytometry validation of RNA-seq targets</u>

A) Representative flow cytometry histograms. B) Graphical analysis of expression of the indicated markers by gated splenic T_R_ populations. Significance determined by one way ANOVA with Tukey’s post-test for pairwise comparisons, ^*^p <0.05, ^**^p < 0.01, ^****^p < 0.001

